# Decreased Interfacial Dynamics Caused by the N501Y Mutation in the SARS-CoV-2 S1 Spike:ACE2 Complex

**DOI:** 10.1101/2021.01.07.425307

**Authors:** Wesam S Ahmed, Angelin M Philip, Kabir H Biswas

## Abstract

Corona Virus Disease of 2019 (COVID-19) caused by Severe Acute Respiratory Syndrome Corona Virus 2 (SARS-CoV-2) has caused a massive health crisis across the globe, with some genetic variants gaining enhanced infectivity and competitive fitness, and thus significantly aggravating the global health concern. In this regard, the recent SARS-CoV-2 alpha variant, B.1.1.7 lineage, reported from the United Kingdom (UK), is of great significance in that it contains several mutations that increase its infection and transmission rates as evident from clinical reports. Specifically, the N501Y mutation in the SARS-CoV-2 spike S1 receptor binding domain (S1-RBD) has been shown to possess an increased affinity for ACE2, although the basis for this is not entirely clear yet. Here, we dissect the mechanism underlying the increased affinity using molecular dynamics (MD) simulations of the available ACE2-S1-RBD complex structure (6M0J) and show a prolonged and stable interfacial interaction of the N501Y mutant S1-RBD with ACE2 compared to the wild type S1-RBD. Additionally, we find that the N501Y mutant S1-RBD displays altered dynamics that likely aids in its enhanced interaction with ACE2. By elucidating a mechanistic basis for the increased affinity of the N501Y mutant S1-RBD for ACE2, we believe that the results presented here will aid in developing therapeutic strategies against SARS-CoV-2 including designing drugs targeting the ACE2-S1-RBD interaction.

**Significance:** The emergence of the new SARS-CoV-2 lineage in the UK in December 2020 has further aggravated the COVID-19 pandemic due to an increased ability of the variant to infect human hosts, likely due to mutations in the viral S1 spike protein including the N501Y S1-RBD mutation that is located at the interface of S1-RBD and ACE2, the host cell receptor for SARS-CoV-2. Given its location at the interface, N501Y S1-RBD mutation can therefore potentially alter the interfacial interaction. Multiple, all-atom, explicit solvent MD simulations of the ACE2-S1-RBD complex carried here indicated a more stable interaction between the N501Y mutant S1-RBD and ACE2 through stabilizing interfacial interactions of residues at one end of the interface that are either sequentially or physically near the mutation site. These mechanistic details will aid in better understanding the mechanism by which the alpha variant has increased infectivity as well as in designing better therapeutics including ACE2-S1 spike protein inhibitors that will, in turn, help thwarting the current and future pandemic.

**Highlights:**

- N501 in the wild type SARS-CoV-2 S1-RBD forms unsustained hydrogen bonds with residues in the ACE2, namely Y41 and K353
- Y501 in the N501Y mutant SARS-CoV-2 S1-RBD is not capable of forming substantial hydrogen bonds with ACE2 within the time span of the current simulation
- Evidence from analyzing the simulation results suggests that Y501 of S1-RBD could form other types of non-covalent interactions with ACE2, such as van der Waals interactions
- N501Y S1-RBD mutation stabilizes the position of interfacial residues neighboring to the mutation site, as well as other non-interfacial residues that are distant from the mutation site
- These altered dynamics results in more stable interaction of S1-RBD with ACE2 which could be the main reason underlying the reported enhanced affinity of S1-RBD in the SARS-CoV-2 alpha variant (UK B.1.1.7 lineage) to ACE2

## Introduction

Severe acute respiratory syndrome coronavirus 2 (SARS-CoV-2) is a positive-sense, single stranded, enveloped RNA virus that belongs to the *Coronaviridae* family, and is the causative agent of Corona virus disease of 2019 (COVID-19).[1] As of August 2021, more than 230 million confirmed cases have been reported worldwide, with more than 4.7 million deaths (September 2021) (https://covid19.who.int/). In general, coronaviruses express four structural proteins: nucleocapsid (N) protein that encapsulates the genomic material; membrane (M) protein that promotes the membrane curvature to bind to the N protein; envelope (E) protein which ensures virus’ assembly and release; and envelope-anchored spike (S) glycoprotein that protrudes from the viral surface and facilitates viral attachment and entry into host cells.[2, 3] The latter is cleaved during viral entry into two subunits, namely S1 and S2.[4] Viral attachment to host cells occurs through binding of its receptor binding domain (RBD) - which is part of the S1 subunit – to the host cell membrane-localized angiotensin converting enzyme 2 (ACE2) receptor. It is important to note that the affinity of SARS-CoV-2 S1-RBD for ACE2 was reported to be 10 times higher than that of SARS-CoV-1, providing a biochemical basis for the increased infection efficiency of SARS-CoV-2 compared to SAR-CoV-1.[5] Indeed, the ACE2-S1-RBD interaction has become an attractive target for inhibiting viral entry into the host cell.[5–9] For instance, the human recombinant soluble ACE2 protein has been utilized for reducing SARS-CoV-2 binding to the cellular ACE2 receptor leading to reduced injury to multiple organs, including the lungs, kidneys, and heart.[10] Similarly, monoclonal antibodies such as 18F3 and 7B11 have been developed to neutralize SARS-CoV-2 infection by blocking epitopes on the S1-RBD.[11]

On top of the increased affinity of SARS-CoV-2 S1-RBD to ACE2 compared to SARS-CoV-1, new genetic variants with increased infectivity and virulence, likely arising under increased immunological pressure in patients suffering from COVID-19 or convalescent plasma therapy [12, 13], have further complicated our efforts towards thwarting the pandemic. One of the key examples of such variants is the S1-RBD D614G mutant that has outcompeted the Wuhan-Hu-1.[14–17] A comparative study conducted by Hou *et al* observed that this variant is superior in infecting the epithelial cells and replicates in higher number than the ancestral virus. The structural analysis showed that the S1-RBD containing the D614G mutation is more flexible and explores the open conformation more than the wild type protein, thus, leading to an increased affinity for ACE2.[15, 18, 19]

Recently, a new phylogenetic group of SARS-CoV-2 (lineage B.1.1.7, alpha variant) has been identified in the COVID-19 Genomics UK Consortium dataset with greater than 50% of the cases belonging to this new cluster (alpha variant) that has an estimated of 50 to 70% increased transmissibility, as per epidemiological and virological investigations.[20, 21] Indeed, reports of the presence of this variant has emerged from other countries as well. Sequence analysis indicates the presence of a total of 17 mutations spanning the ORF1ab, spike, Orf8 and the N protein in the genome of this variant.[21] Majority of these mutations (8 out of the total 17), however, are present in the spike protein. These include deletion mutations (ΔH69V70 and ΔY144) and missense mutations (N501Y, A570D, P681H, T716I, S982A and D1118H). Of these, the N501Y substitution strikes out as one of the most interesting mutations due to its presence at the ACE2-S1-RBD interaction interface [22], raising the possibility of an altered interaction between the two proteins. In fact, deep mutational analysis of S1-RBD, in combination with the yeast-surface-display platform, has revealed an increased affinity of the N501Y mutant S1-RBD to ACE2 (apparent *K*_d_ of 3.9×10^−11^ M for the wild type vs. 2.2×10^−11^ M for the N501Y mutant).[23] In addition to that, a recent study demonstrated the EC_50_ of the mutant RBD (possess nine mutations I358F, V445K, N460K, I468T, T470M, S477N, E484K, Q498R, N501Y) was nearly 17 times lower than that of the RBD-WT.[24] Further, these serve as an evidence for the constantly evolving SARS-CoV-2 with more contagious mutations spreading rapidly.

In the current study, we performed multiple all atom, explicit solvent MD simulations to gain an insight into the mechanism underlying the increased affinity of the N501Y mutant S1-RBD for ACE2. Simulations of the wild type and the N501Y mutant S1-RBD in complex with ACE2 showed a prolonged and stable interaction between the Y501 residue with the neighbouring Y41 and K353 residues in ACE2 in the mutant complex as compared to the N501 residue in the wild-type complex. Importantly, these simulations also revealed a localized decreased dynamics for interfacial residues in the mutant as compared to the wild-type complex that led to changes in interfacial interactions of these residues. Although these were most noticeable for residues near the N501Y S1-RBD mutation site, stabilized dynamics were also observed for non-interfacial positions that are distant from the N501Y S1-RBD mutation site.

## Materials & Methods

### ACE2-S1-RBD structure preparation

The three-dimensional structure of ACE2-S1-RBD complex spanning residues S19 to D615 of ACE2 and T333 to G526 of spike (S) glycoprotein was obtained from the RCSB PDB database as a PDB file (PDB ID: 6M0J).[22] PyMOL (The PyMOL Molecular Graphics System, Version 2.0.0, Schrödinger, LLC; https://pymol.org) was used to visualize the three-dimensional structure and to generate the N501Y mutant structure using the Mutagenesis tool available in PyMOL. Wild-type and mutant PDB structure files were exported after removing ions and solvent molecules.

### ACE2-S1-RBD molecular dynamics simulations

Molecular dynamics simulations were performed using NAMD V2.13 software [25] and CHARMM36 force field [26], as described previously [27]. The simulation system consisting of the biomolecular complex formed by the ACE2-S1-RBD was generated from the previously prepared PDB files using the QwikMD Toolkit [28] available as a plugin in Visual Molecular Dynamics (VMD) [29] software V1.9.3. Briefly, the proteins were solvated using TIP3P (transferable intermolecular potential with 3 points)[30] cubic water box and charges were neutralized using 0.15 M NaCl final concentration in explicit solvent with Periodic Boundary Conditions applied. The biomolecular simulation systems consisted of ~ 453000 atoms. Energy minimization was first performed for 1000 timesteps, followed by a thermalization step where the system was slowly heated for 0.25 ns using a temperature ramp where the temperature was raised from 60 K to 310 K at 1 K increment. Temperature was then maintained at 310 using Langevin temperature control and at 1.0 atm using Nose-Hoover Langevin piston control and a 1 ns constrained equilibration step was then performed where protein backbone atoms where constrained using harmonic potential. Finally, 50 ns duplicate runs were performed for each of the WT and Mutant complex systems. A 2 fs time step of integration was chosen for all simulation where short-range non-bonded interactions were handled at 12 Å cut-off with 10 Å switching distance, while Particle-mesh Ewald (PME) scheme was used to handle long-range electrostatic interactions at 1 Å PME grid spacing. Trajectory frames were saved every 10,000 steps.

### ACE2-S1-RBD molecular dynamics simulation trajectory analysis

Analysis of the trajectories was performed using the available tools in VMD software.[29] Independent root-mean-square deviation (RMSD) calculations of backbone atoms of ACE2 and S1-RBD proteins were performed using the “RMSD trajectory Tool” in VMD.[29] Root-mean-square fluctuations (RMSF) measurements were performed for protein Cα atoms. Hydrogen bonds analysis between ACE2 and S1-RBD was performed at a cut-off distance of 3.5 Å and a cut-off A-D-H angle of 20° using the “Hydrogen Bonds” analysis extension in VMD [31, 32]. Interfacial residues were determined from the available ACE2-S1-RBD complex (PDB ID: 6m0j) at a cut-off distance of 5 Å using PyMOL. Energy calculations were performed using “NAMD Energy” analysis tool available as part of VMD. Center of mass distances between paired selections were determined using VMD. Standard deviations of the inter-residue distances obtained over the course of the simulation were then normalized with their respective initial distances and plotted as a ratio of N501Y mutant to WT ACE2-S1-RBD complexes. Dynamic Cross-Correlation (DCC) analysis was performed using the DCC algorithm from MD-TASK software suite [33] for analyzing molecular dynamics trajectories (https://mdmtaskweb.rubi.ru.ac.za/) as well as by using Bio3D R package [34, 35]. DCC calculations were based on the position of Cα atoms obtained after aligning trajectory frames on the Cα atoms of the original complex structure. Average DCC figures were prepared using MATLAB and results were represented as heat maps that indicate the range of correlations from +1 (high correlation) to 0 (no correlation) to −1 (high anti-correlation). All analysis were performed based on 500 trajectory frames (10 frames/ns).

The representative composite timestep snapshot images were prepared by saving the trajectory coordinates as PDB file format every 5 ns and then combining a total of 11 frames to form the composite images. Representative trajectory movies of the 50 ns simulations were prepared from 500 trajectory snapshots (10 snapshots/ns) generated using VMD Movie Maker Tool [29] and compiled using Fiji distribution of ImageJ software [36] at a frame rate of 30 fps.

### Data Analysis and Figure Preparation

GraphPad Prism (version 9 for macOS, GraphPad Software, La Jolla California USA; https://www.graphpad.com), in combination with Microsoft Excel, were used for data analysis and graph preparation. Figures were assembled using Adobe Illustrator.

## Results & Discussion

In order to understand the mechanism underlying the enhanced affinity of the N501Y mutant over the wild type S1-RBD for ACE2, we initiated MD simulations with the available ACE2-S1-RBD complex structure (PDB ID: 6M0J)[22] (Fig. 1). A closer inspection of the ACE2-S1-RBD interface indicated that residues Y41 and K353 of ACE2 are in close proximity to the N501 residue of S1-RBD (Fig. 1; outset). In fact, N501 has been reported to participate in hydrogen bonding (at 3.7 Å distance) with Y41 residue of ACE2, indicating its potential role in the ACE2-S1-RBD interaction.[22] We hypothesized that this interaction at residue-level is altered by the N501Y mutation of S1-RBD. We also hypothesized that other pair-wise interactions at the interface may be altered by the same mutation. To test these hypotheses, we initiated multiple, all-atom MD simulations in explicit solvent with the wild type and the N501Y mutant ACE2-S1-RBD complex structure and analyzed the trajectories obtained for general structural dynamics and specific interactions. Further, we performed the simulations in duplicates to test the consistency of the results and for statistical support.

**Figure 1.**
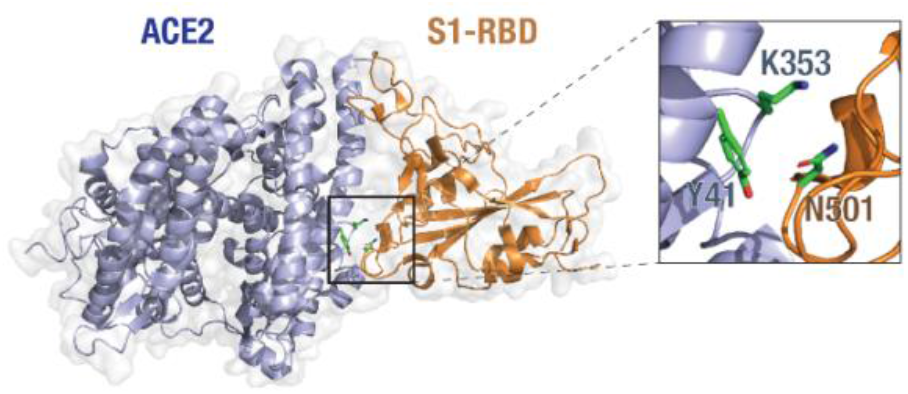
ACE2-S1-RBD complex and interactions of the N501 residue. Cartoon representation of the ACE2-S1-RBD structure (PDB: 6M0J[22] showing the relative positioning of residues Y41 and K353 in ACE2 (light blue) and residue N501 in S1-RBD (orange).

These MD simulations revealed a generally decreased dynamics of the N501Y mutant ACE2-S1-RBD complex compared to the wild type complex as seen from the composite image of the complexes obtained from the simulation trajectories (Figure 2A).[37–43] However, RMSD analysis of backbone atoms of the proteins ACE2 and S1-RBD individually, taken over the entire course of simulation, did not show any clearly discernable trend for structural evolution of amino acid residues in the complex (Figure 2B,C). This suggests that any alteration in the biochemical interaction between the two proteins likely arises due to changes in the dynamics of specific, individual residues in the proteins. Indeed, RMSF analysis of individual amino acid residues in the proteins showed several distinct changes, with a general decrease in the N501Y mutant complex (Figure 2D). Specifically, in ACE2, residue positions at the N-terminal (from 19 until 111), central (183 until 206) and to a smaller extent at the C-terminal (from 542 until 588) of ACE2 showed a reduced RMSF values in the N501Y mutant complex. Importantly, reduced RMSF values were observed for the Y41 and K353 residues in ACE2 in the mutant complex. On the other hand, residues 281 to 283 in ACE2 showed an increased RMSF value in the mutant complex. RMSF analysis of S1-RBD showed a reduced structural fluctuation of Y501 in the mutant complex compared to N501 in the wild-type complex (Figure 2E), indicating a more stable interaction with adjacent, interfacial residues in ACE2. In addition, residue positions from 362 until 395 showed a substantially reduced RMSF values in the mutant complex (Figure 2E). The latter is suggestive of the possibility of an allosteric effect of the N501Y S1-RBD mutation on the mutant ACE2-S1-RBD complex as compared the wild-type complex.[37–43]

**Figure 2.**
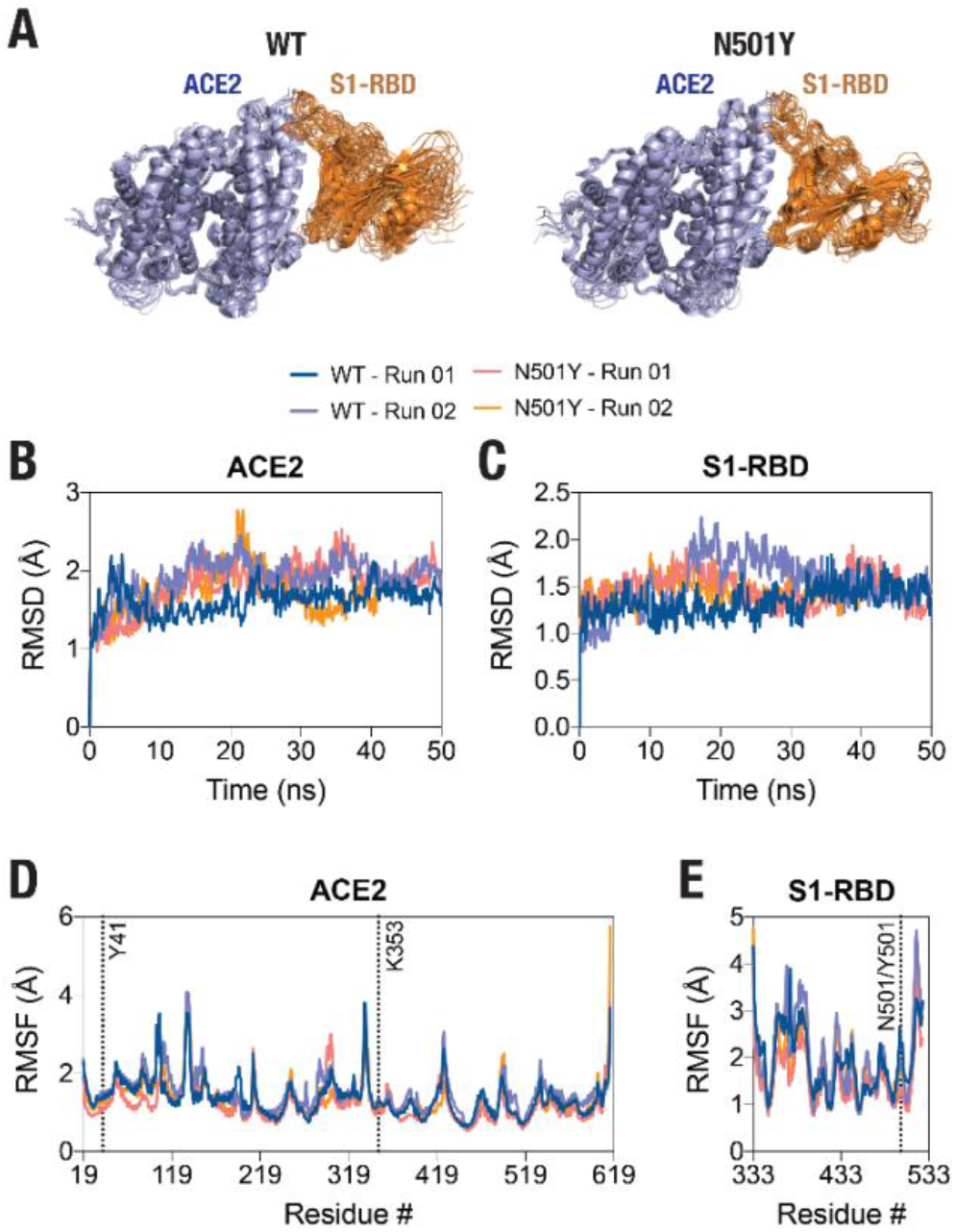
Decreased structural dynamics of the N501Y mutant S1-RBD in complex with ACE2. (A) Cartoon representation of the wild type (left panel) and the N501Y mutant (right panel) ACE2-S1-RBD complex showing structural evolution of the complex over time in a 50 ns all-atom, explicit solvent MD simulation. Composite images were prepared using 11 consecutive frames from up to 50 nm simulations with each frame 5 ns apart. (B,C) Graph showing backbone (Cα) root-mean-square deviation (RMSD) values of ACE2 (B) and S1-RBD (C) obtained from the simulation of the WT and N501Y mutant ACE2-S1-RBD complexes. (D,E) Graph showing backbone (Cα) root-mean-square fluctuation (RMSF) values of ACE2 (D) and S1-RBD (E) obtained from up to 50 nm simulations of the WT and N501Y mutant ACE2-S1-RBD complexes.

Following these analyses, we determined the residue-residue distances based on the center-of-mass between position 501 in S1-RBD and key residues, Y41 and K353, in ACE2 of the ACE2-S1-RBD complexes, as they evolve during the span of the simulations (Figure 3A). First, N501 residue in the wild-type complex showed a substantially higher structural fluctuations in comparison to Y501 in the mutant complex. In fact, as the simulation progressed, N501 in the wild type S1-RBD tended to move away from the ACE2-S1-RBD interface, with ACE2 Y41 residue moving in the other direction in the first simulation run (Figure 3A; left panel, Supporting Movie 1). This was not the case for N501Y S1-RBD mutant, in which Y501 sustained its contact at the ACE2-S1-RBD interface over the entire simulation time (Figure 3A; right panel, Supporting Movies 3 and 4). Indeed, the inter-residue distance analysis revealed a dramatic increase in the distance between Y41 and K353 in ACE2 and N501 in S1-RBD after about 30 ns in the first simulation run while a smaller, more fluctuating, increases at different times were seen in the second run (Figure 2B, C). This is in contrast to the distances measured for the same pair of ACE2 residues with Y501 in the mutant complex (~7 and ~4.5 Å, respectively) (Figure 2B, C). These data suggests that Y501 residue of N501Y mutant S1-RBD forms more stable interactions at the interface with Y41 and K353 residues of ACE2 compared to the WT. To determine if the N501Y mutation impacts interaction at the opposite end of the ACE2-S1-RBD interface, we monitored the inter-residue distances between the hydrogen bond-forming Q24 of ACE2 and N487 of S1-RBD and the closely juxtaposed (but not in hydrogen bond) T27 in ACE2 and Y489 in S1-RBD [22]. In contrast to the observations made with the Y41-N501 and K353-N501 pairs, these pairs did not show substantial difference in fluctuations of their relative positioning (Figure 3D, E) compared to the mutant complex, suggesting that the effect of the N501Y mutation on the interface may be local, and does not affect the inter-chain interaction of ACE2 and S1-RBD interface in the timescales that we have explored here.

**Figure 3.**
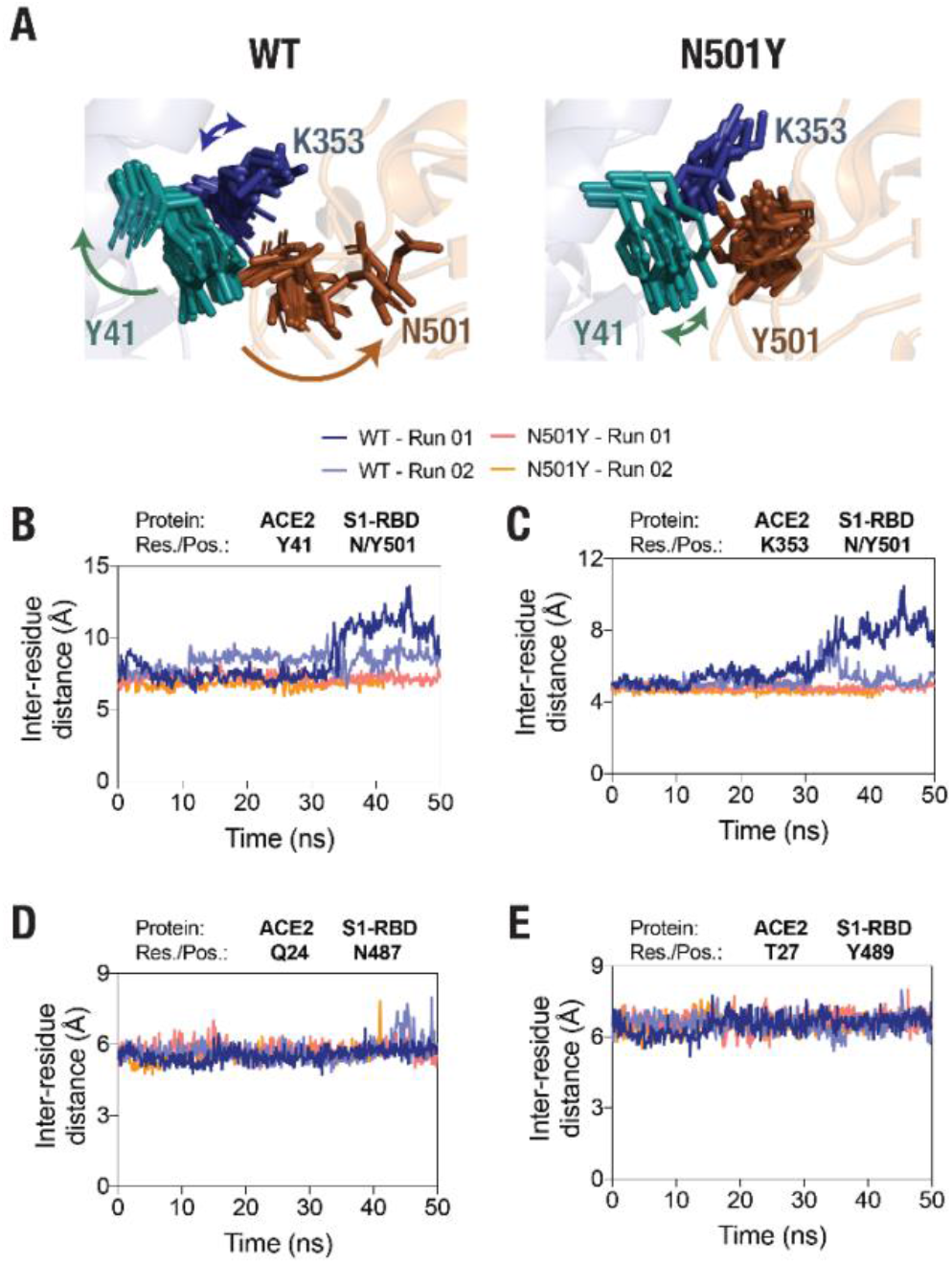
Sustained interaction of S1-RBD Y501 residue (N501Y mutant) with ACE2. (A) Temporal evolution of residues Y41 and K353 in ACE2 and either the N501 in the WT S1-RBD (left panel) or the Y501 in the N501Y mutant S1-RBD (right panel) in the MD simulation. A total of 11 frames obtained from up to 50 nm simulations, each 5 ns apart, were compiled together. Note the increased fluctuation of the N501 residue in the wild type S1-RBD. (B,C,D,E) Graph showing inter-residue distances between the center of masses of residue Y41 in ACE2 and N501 in the wild type and Y501 in the N501Y mutant S1-RBD (B), K353 in ACE2 and N501 in the wild type and Y501 in the N501Y mutant S1-RBD (C), Q24 in ACE2 and N487 in either the wild type or the N501Y mutant S1-RBD (D), and T27 in ACE2 and Y489 in either the wild type or the N501Y mutant S1-RBD (E). Note the increased inter-residue distance between the residues Y41 and K353 in ACE2 and N501 in S1-RBD in the wild type ACE2-S1-RBD complex (B,C) compared to the N501Y mutant complex.

We then attempted to determine if there are any correlated confirmational dynamics of the complex in the wild type and the N501Y mutant using dynamic cross-correlation (DCC) analysis, which is used to detect correlated motions in protein segments. Application of a minimum cut-off of 0.8 to positive DCC values obtained from individual MD runs showed a generally greater correlated motions (both positive as well as negative) in the wild type ACE2-S1-RBD complex compared to the N501Y mutant complex. However, DCC analysis did not reveal any dynamically correlated motions between N501 of S1-RBD or any other interfacial residues located near this position and residues in ACE2 in the WT complex. Although, in the S1-RBD mutant complex, high dynamical cross-correlations were observed between residues Y501 and G502 of S1-RBD on one side and ACE2 interfacial residues, namely K353, G354, and D355, on the other side (Figure 4A). Interestingly, application of the same cut-off to the negative DCC values revealed a higher anti-correlated motions between the two chains in the WT complex compared to the mutant complex (Figure 4A). Moreover, by averaging the DCC values for the two runs, our results revealed higher dynamical cross-correlated motions between cluster of interfacial residues sequentially adjacent to the mutation site in the N501Y mutant S1-RBD (residues G496, Q498, T500, Y501, G502, V503, Y505) on one side and ACE2 interfacial clustered positions (S19, Q24, T27, F28, D30, K31, H34, E35, E37, D38, Y41, Q42, L45), (Q325, G326, N330), and (A386, R393) on the other side, compared to the WT ACE2-S1-RBD complex (Figure 4B). Similar observations were made for the DCC values between all the aforementioned ACE2 clustered positions and S1-RBD clustered residues (V445, G446, and Y449) which are physically adjacent to the mutation site as they are located on the same end of the interface as the N/Y501 clustered position mentioned earlier. Additionally, the average DCC analysis revealed a global decrease in the significantly dynamic anti-correlated motions in the mutant compared to the WT complex (Figure 4B). These results provide insight on the effect of the N501Y mutation on the dynamics of interfacial residues adjacent, either in protein sequence or in terms of physical location, to the mutation site and the distant effect of the mutation on the dynamics of non-interfacial residues manifested as a decrease in the anti-correlated inter-chain motions in the mutant complex. The decrease in the inter-chain dynamic anti-correlations could probably be due to a non-interfacial, distant stabilizing effect of the mutation as indicated by here by the RMSF analysis.

**Figure 4.**
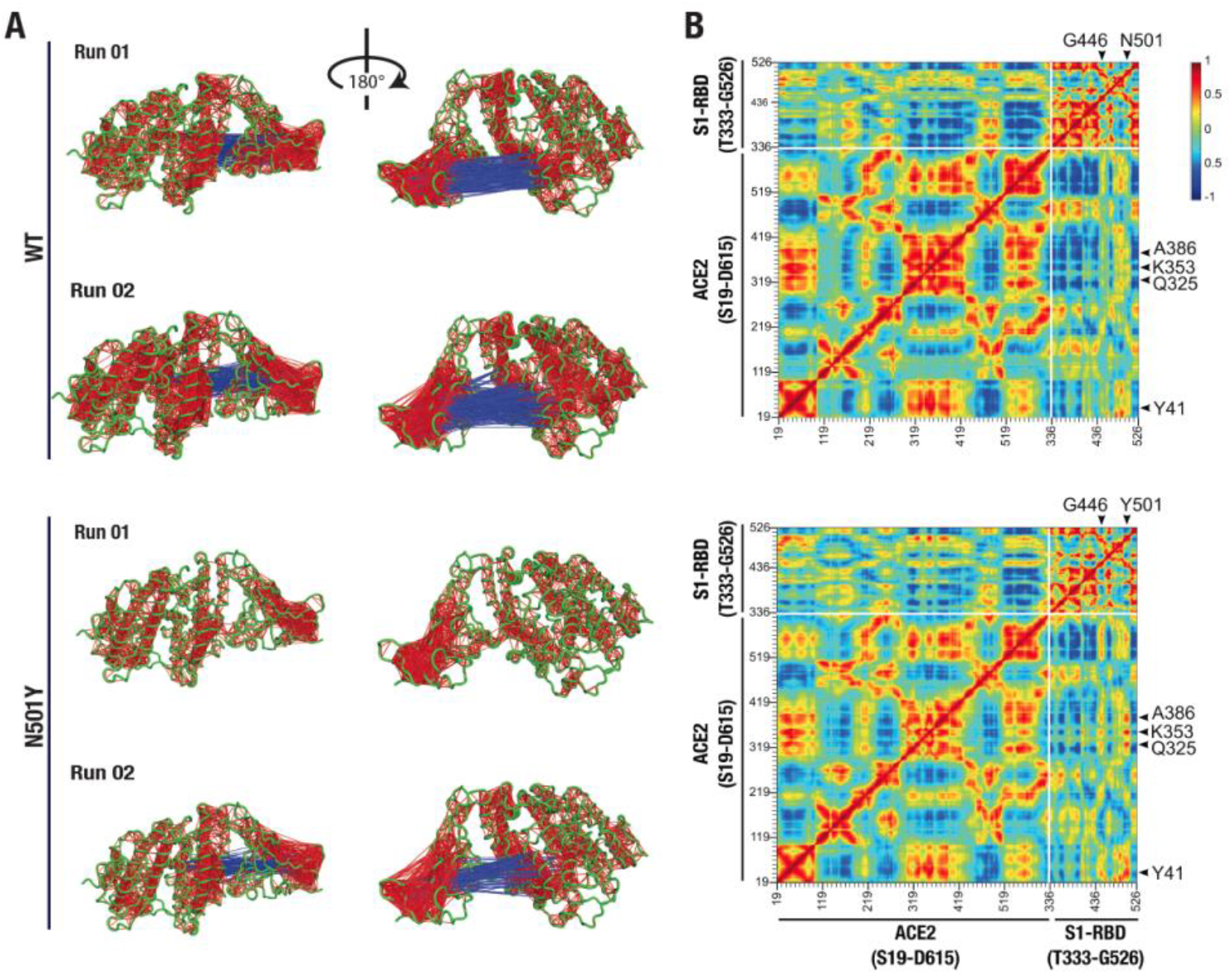
Altered dynamical cross-correlated motions in the ACE2-S1-RBD N501Y mutant complex. (A) Cartoon representation of ACE2-S1-RBD wild type (top two panels) and N501Y mutant (bottom two panels) complex showing DCC values (cut-off, ±0.8). Note the positively correlated motions observed between Y501 and G502 on S1-RBD and residues K353, G354, and D355 on ACE2 in the mutant complex but not in the wild type complex while less dynamically anti-correlated motions were observed in the mutant complex compared to the wild type complex. (B) Heat map showing average DCC values from two independent 50 ns MD simulations of the wild type (top panel) and the N501Y mutant ACE2-S1-RBD complex (cut-off, ±0.8). Note the higher inter-chain dynamically cross correlated motions between residues at the interface. Also note the global decrease in the anti-correlated motions in the mutant complex.

In order to better understand how the two proteins interact at the interface and how this interaction compares in the WT and mutant complexes, we next performed interfacial hydrogen bond formation occupancy time analysis using a 3.5 Å cut-off distance and 20° cut-off angle. By applying a cut-off trajectory occupancy time of 5%, we were able to identify 22 unique hydrogen bonds that form at the interface during the span of the simulation time by either the main chain or side chain of residues (Figure 5A). Interestingly, this analysis revealed that position 501 of S1-RBD is capable of hydrogen bond formation with residues Y41 and K353 of ACE2 in the WT complex but not in the mutant complex. In fact, Y501 in the S1-RBD mutant complex did not form any substantial H-bonds with residues in ACE2. This indicates that Y501 residue in the mutant S1-RBD does not contribute to significant hydrogen bond formation at the interface, but rather may be involved in forming other types of noncovalent interactions. In fact, by calculating interaction energy between this position and interfacial residues in ACE2, we found that this position forms additional, and more sustained, van der Waals interactions at the interface (Supporting Figure 1). These results are in contrast with previous reports suggesting enhanced H-bond formation by Y501 in the mutant complex [21, 44, 45] driving the enhanced binding affinity of N501Y S1-RBD mutant to ACE2 [46–48]. More importantly, by calculating the difference between mean % occupancy time, we were able to determine changes in the % occupancy time between hydrogen bonds formed in WT and mutant complexes. Interestingly, residues immediately adjacent to the 501 position in S1-RBD (T500 and G502) had the highest change in the H-bond occupancy (+36.5% and +22.5%, respectively), further indicating that the local effect of the mutation on the interface. A closer inspection of the distances – from a snapshot taken 40 ns in the simulation time - between residues that possessed the highest increase (ACE2-K353-mainchain:S1-RBD-G502-mainchain, ACE2-D355-sidechain:S1-RBD-T500-sidechain, and ACE2-D30-sidechain:S1-RBD-K417-sidechain) and decrease (ACE2-K353-sidechain:S1-RBD-Q498-sidechain, ACE2-K353-sidechain:S1-RBD-496,-mainchain and ACE2-D38-sidechain:S1-RBD-Y449-sidechain) in the % mean occupancy time, revealed that the distances between the interacting selections was decreased and increased, respectively, in the mutant complex compared to the WT complex (Figure 5B). More importantly, distance measurements revealed that in both cases (increased and decreased hydrogen bond mean occupancy time) there was much less distance fluctuations between H-bond forming residue pairs in the mutant complex compared to the WT. Interestingly, pair-wise distances that involved S1-RBD hydrogen bond forming residues at positions that sequentially distant, but physically adjacent to the mutation site, were also less fluctuating in both cases of increased (ACE2-D30-sidechain: S1-RBD-K417-sidechain) and decreased (ACE2-D38-sidechain:S1-RBD-Y449-sidechain) hydrogen bond formation % mean occupancy time (Supporting Figure 2). However, this distant effect did not affect key H-bond forming interfacial residues that are distant and physically non-adjacent to the mutation site. For instance, when looking at the distance between interfacial residues that contribute to substantial hydrogen bonding at the interface, but yet have a mean occupancy time not changing with the mutation (namely ACE2-Y83-sidechain:S1-RBD-N487-sidechain, and ACE2-E35-sidechain:S1-RBD-Q493-sidechain), we could see that these residues are not located near the mutation site and have no marked difference in distance fluctuation between the wild type and mutant complexes (Supporting Figure 3A), indicating that interfacial residues that are sequentially or physically adjacent to the mutation site are most affected by the mutation. The same can be concluded from calculating the distance between close-by interfacial residues at the far opposite end of the interface as was described above (Figure 3D and E.).

**Figure 5.**
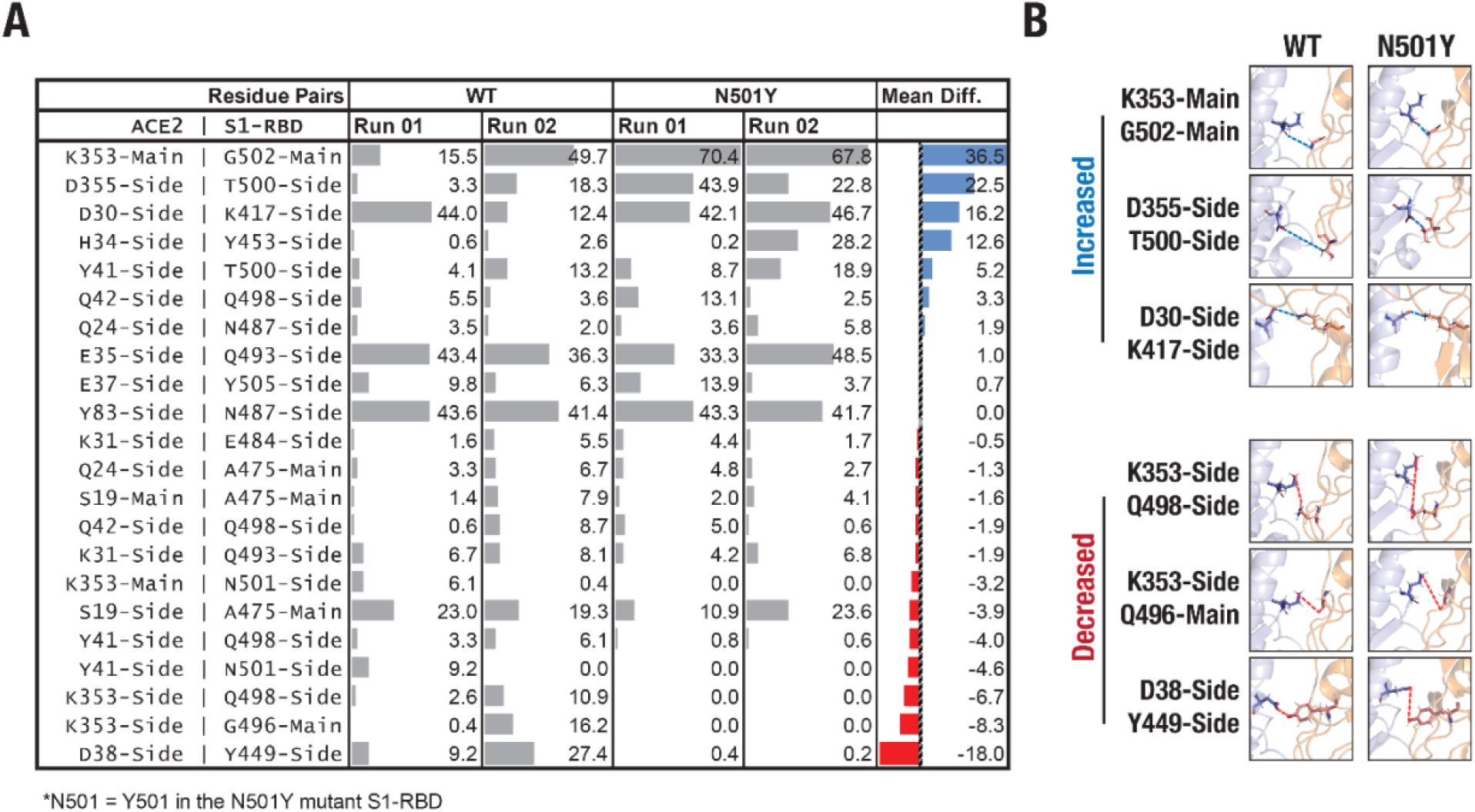
Altered H-bonding between ACE2-S1-RBD interfacial residues. (A) Table showing hydrogen bonds formed at the interface by applying a cut-off of 3.5 Å distance, 20° angle, and ≥ 5% occupancy time. Numbers represent the % occupancy time of the hydrogen bond per the total simulation time. (B) Cartoon representation of selected interfacial residue pairs showing greatest alteration in H-bond interaction. Panels: interfacial residue pairs showing increased (top panel) or decreased (bottom panel) H-bond occupancy inter-residue distance between pairs that display the highest change (increase, blue; decrease, red) in the hydrogen bond occupancy time between wild type and mutant complexes, obtained from a snapshot taken at 40 ns. Mean Diff.: Mean difference, i.e., difference between hydrogen bond mean percent occupancy time of the wild type and mutant complexes. Blue and red colors indicate percent increase and decrease in occupancy time, respectively.

To further confirm the effect of N501Y mutation on the interface, we calculated pair-wise residue distances between residues that form the ACE2-S1-RBD interface. Using a cut-off distance of 5 Å, we were able to identify 25 interfacial residues in ACE2 and 22 in S1-RBD (Figure 6A). Pairing these residue, distance-wise, resulted in 550 pairs. The mean and standard deviation of 500 distance measurements (obtained from 500 trajectory frames) for each pair were then calculated. The average standard deviation was the calculated for all pairs in the wild type and mutant complexes. A heatmap was created by dividing the two-runs average standard deviations of the pair-wise distances in the mutant complex over those in the wild-type complex (Figure 6B). Therefore, a value of higher than 1.0 will indicate a higher pair-wise distance fluctuation in the mutant complex compared to the wild type, while the opposite is true for values less than 1.0. Overall, this analysis revealed a general stabilizing effect of the mutation on residues at the interface, as can be inferred from the lowest and highest values of the heatmap (0.4 vs 1.2). In fact, this stabilizing effect was more prominent on residues that are adjacent to the mutation site either sequentially (T500, N501, G502, V503, and Y505) or physically (V445, G446, and Y449), which further supports our findings in this study on the local stabilizing effect of the mutation on residues at the interface (Figure 5C). Interestingly, the heatmap showed that pair-wise distances that involve T500 were the most stabilized by the mutation, rather than pair-wise distances that involve the mutation site itself (Figure 5B).

**Figure 6.**
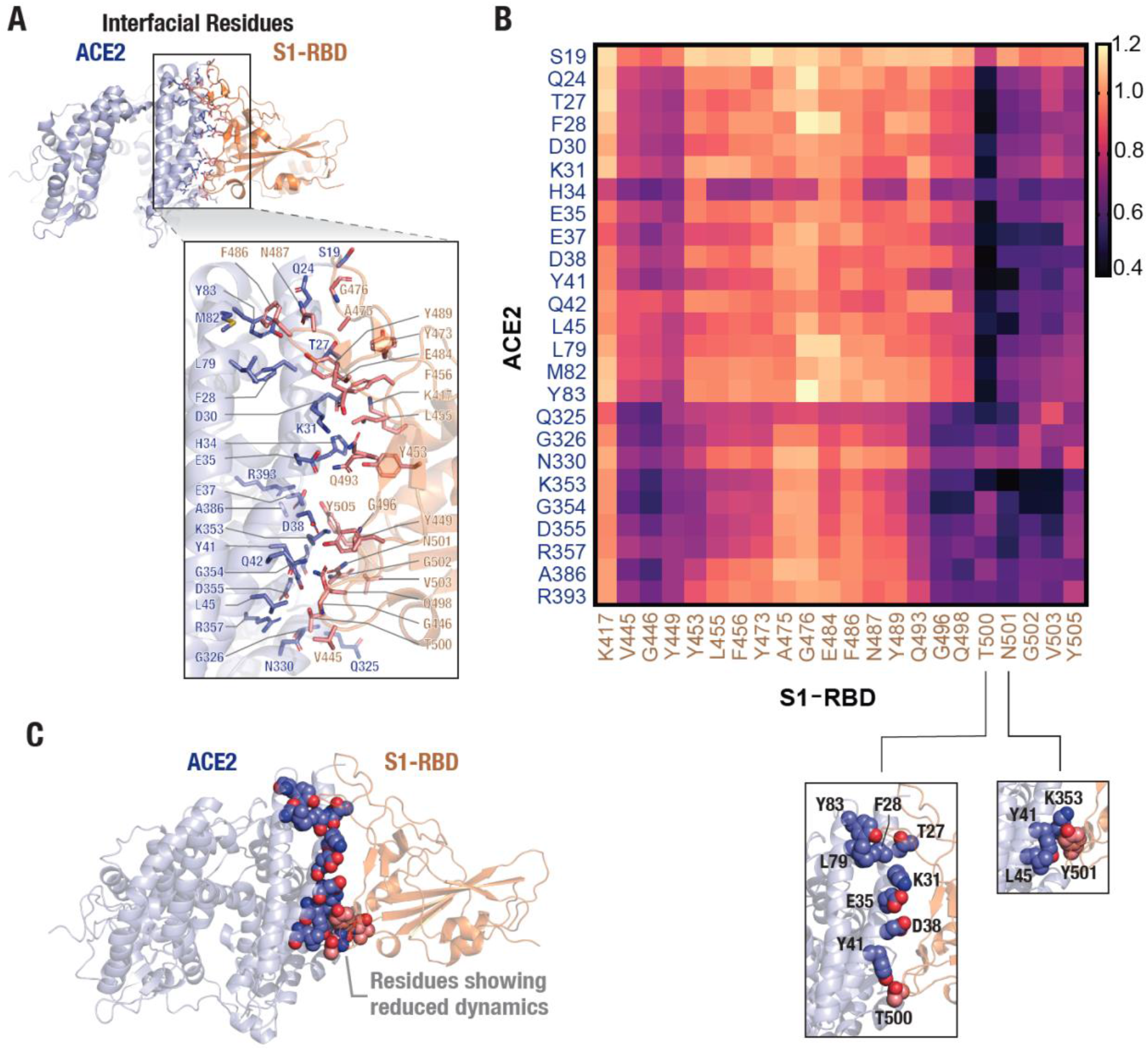
Stabilizing effect of the N501Y mutation on the ACE2 and S1-RBD interfacial interaction. (A) Cartoon representation showing interfacial residues determined from the available ACE2-S1-RBD complex (PDB ID: 6m0j) at a cut-off distance of 5Å. A total of 25 residues (blue) and 22 residues (orange) were identified at the interface in ACE2 and S1-RBD, respectively. (B) Heatmap representing ratio of mean standard deviation of inter-residue distances (normalized to initial distances) in the N501Y and WT ACE2-S1-RBD. Note the decrease in the dynamics of Y501 and residues neighboring the mutation site, indicating a stabilizing effect of the mutation. (C) Schematic showing reduced dynamics of interfacial residues in the N501Y mutant. ACE2 (blue) and S1-RBD (brown) complex showing interfacial residues with reduced dynamics in the N501Y mutant in comparison to the wild type interface.

## Conclusion

To conclude, the MD simulations performed here with the ACE2-S1-RBD complex provide an unambiguous mechanistic insight into the increased binding affinity of the N501Y mutant S1-RBD for ACE2. Specifically, our computational work shows that the mutation of N501 residue into a Y results in a stable interaction with the Y41 and K353 residues in ACE2. This is positively impacted by the altered dynamics of the S1-RBD upon N501Y mutation, which is more noticeable on residues adjacent to mutation site, and extends to include certain nonadjacent residues, although the reason behind it is not entirely clear and will likely require further investigation. Even though experiments determining binding of fluorescently labelled ACE2 and S1-RBD displayed on yeast cells and computational results presented here clearly indicate an increased affinity, it remains to be seen if the N501Y mutation alone can increase the overall fitness of the virus. The N501Y and associated mutations in the S1 spike protein has gained tremendous interest of the scientific community given that this lineage of SARS-CoV-2 has been suggested to be behind the dramatic increase in the number of COVID-19 cases in UK. We believe that the results outlined here will be helpful in efforts towards thwarting this new wave of COVID-19 by enabling discovery of potent inhibitors of ACE2-S1-RBD interaction [5–9].

## Supporting information

MD trajectory movie of the wild type ACE2-S1-RBD complex

MD trajectory movie of the mutant ACE2-S1-RBD complex

## Acknowledgements

This work is supported by an internal funding from the College of Health & Life Sciences, Hamad Bin Khalifa University, a member of the Qatar Foundation. W.A. and A.M.P. are supported by scholarship from the College of Health & Life Sciences, Hamad Bin Khalifa University, a member of the Qatar Foundation. Some of the computational research work reported in the manuscript were performed using high-performance computer resources and services provided by the Research Computing group in Texas A&M University at Qatar. Research Computing is funded by the Qatar Foundation for Education, Science and Community Development (http://www.qf.org.qa).

## Author Contributions

K.H.B. conceived the experiments. W.S.A, A.M.P. and K.H.B. performed experiments, analyzed data, prepared figures and wrote the manuscript. All authors reviewed and approved the manuscript.

## Competing interests

The authors declare no competing interests.

## Supporting Information

### Supporting Movies

**Supporting Movie 1. Trajectory movie of WT complex first simulation run displaying interaction between ACE2 and WT S1-RBD.** Movie was created by compiling 500 snapshots over 50 ns simulation time (10 snapshots/1ns) using a frame rate of 30 fps.

**Supporting Movie 2. Trajectory movie of WT complex second simulation run displaying interaction between ACE2 and WT S1-RBD.** Movie was created by compiling 500 snapshots over 50 ns simulation time (10 snapshots/1ns) using a frame rate of 30 fps.

**Supporting Movie 3. Trajectory movie of N501Y S1-RBD mutant complex first simulation run displaying interaction between ACE2 and N501Y mutant S1-RBD.** Movie was created by compiling 500 snapshots over 50 ns simulation time (10 snapshots/1ns) using a frame rate of 30 fps.

**Supporting Movie 4. Trajectory movie of N501Y S1-RBD mutant complex second simulation run displaying interaction between ACE2 and N501Y mutant S1-RBD.** Movie was created by compiling 500 snapshots over 50 ns simulation time (10 snapshots/1ns) using a frame rate of 30 fps.

## Supporting Figures

**Supporting Figure 1.**
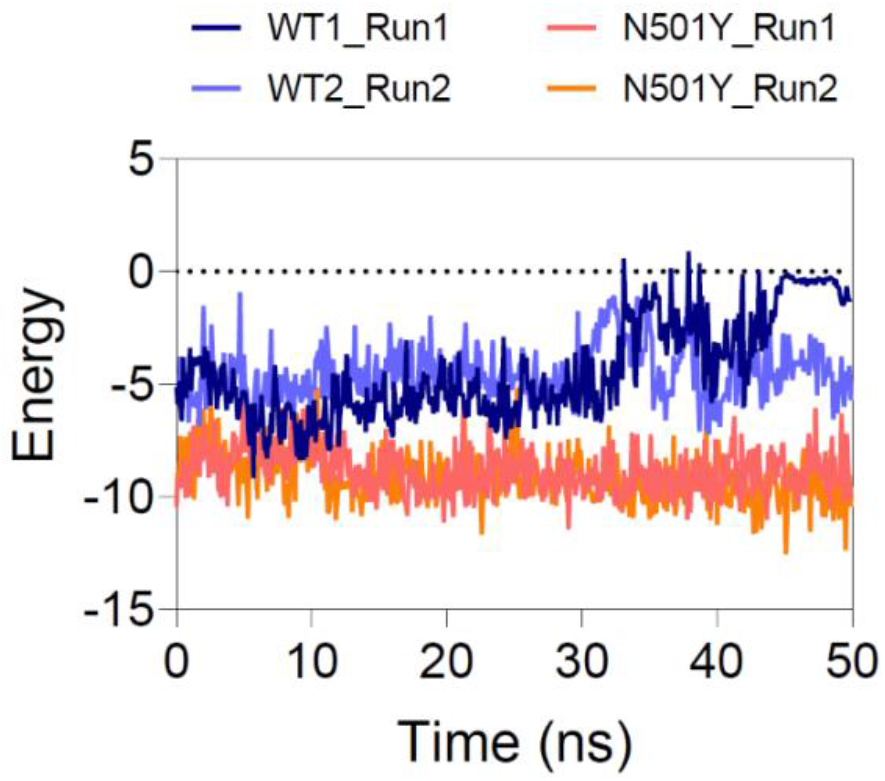
van der Waals energy of interaction between position 501 of S1-RBD and residues in ACE2 in the wild type and mutant complexes.

**Supporting Figure 2.**
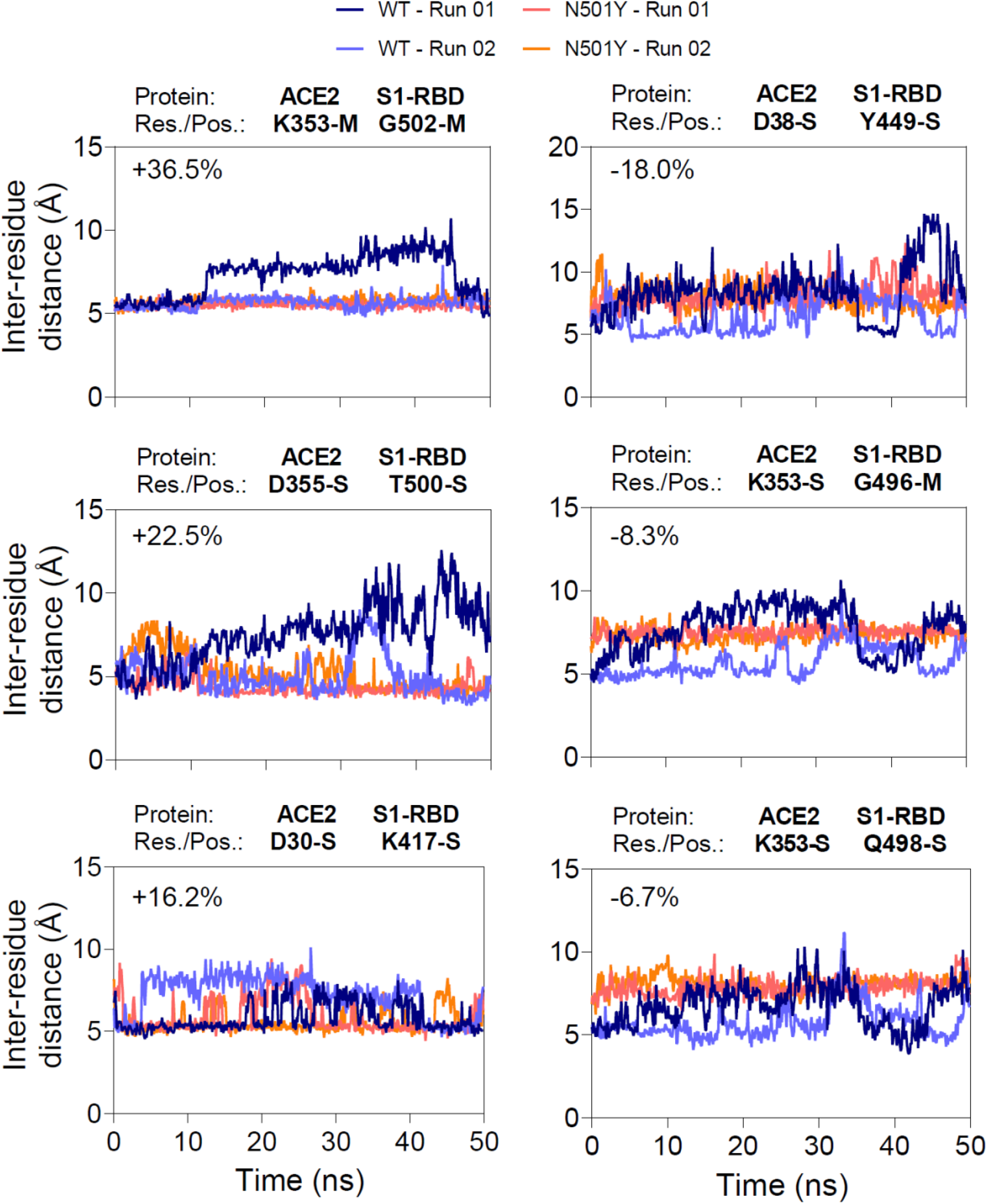
Analysis of distance between interfacial residues whose hydrogen bond formation were most affected by the N501Y S1-RBD mutation. On the left panel, graphs showing inter-selection (mainchain or sidechain) distances between hydrogen bond forming residues that showed the highest increase in the % mean occupancy time, arranged from the highest increase (top) to the lowest increase (bottom). On the right panel, graphs showing inter-selection (mainchain or sidechain) distances between hydrogen bond forming residues that showed the highest decrease in the % mean occupancy time, arranged from the highest decrease (top) to the lowest decrease (bottom). Note the more stable, less fluctuating, hydrogen bonds formed by pairs in the mutant complex. Inset percentages represent the increase (+) and decrease (−) in hydrogen bond mean occupancy time between the wild type complex and N501Y mutant S1-RBD complexes. Res: residue, Pos: position, S: side chain, M: main chain.

**Supporting Figure 3.**
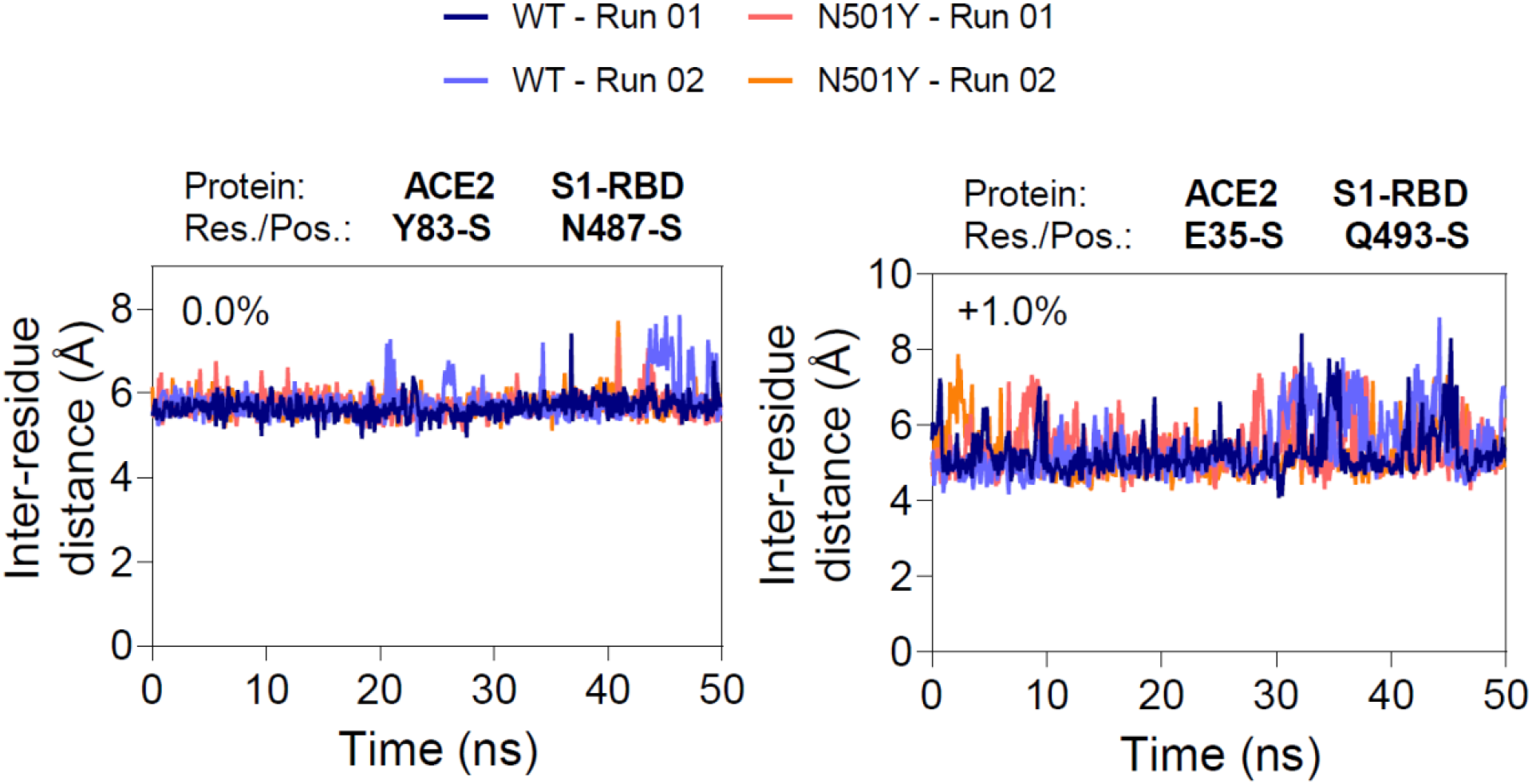
Analysis of distances between residues that form substantial hydrogen bonding at the interface and whose hydrogen bond formation were not affected by the N501Y S1-RBD mutation. Note the inconsiderable difference in the distance fluctuation between pairs in the wild type and mutant complex. Inset percentages represent the percent change in hydrogen bond mean occupancy time between the wild type complex and N501Y mutant S1-RBD complexes. Res: residue, Pos: position, S: side chain.

## Notes

### Competing Interest Statement

The authors have declared no competing interest.

### Summary of Updates

We have included additional analysis and results in the edited manuscript.

